# The ambiguity of “hybrid swarm”: inconsistent definitions and applications in existing research

**DOI:** 10.1101/2024.10.11.617831

**Authors:** Jillian N. Campbell, Elizabeth G. Mandeville, Nathan C. Lewis, Amanda V. Meuser

## Abstract

Hybridization is common in wild taxa and often increases in frequency following anthropogenic disturbance to an environment. Next-generation sequencing techniques make genomic analysis of a large number of individuals feasible, vastly improving the analysis for and promoting a greater frequency of studies on hybridization. However, terminology surrounding hybridization can be inconsistent; in particular, the term “hybrid swarm” has been used extensively in the literature but lacks a consistent definition. In this paper, we conducted a comprehensive review of the literature that uses the term “hybrid swarm” in reference to hybridization between taxa and challenged putative definitions of the term. We found that the term “hybrid swarm” is used in a variety of contexts, including some contradictory to other literature, and that there is little consensus on what constitutes a hybrid swarm in terms of hybrid outcomes, frequency relative to disturbances, or duration of existence. We dissuade researchers from use of the term “hybrid swarm” and instead suggest more specific and clear terminology to describe aspects of hybridization. Consequently, we hope that this paper promotes consensus surrounding hybridization terminology and improves the quality of future research on hybridization.

## 2 Introduction

Hybridization between species can result in outcomes that range from speciation, to introgression of genes, or to production of sterile hybrids (Allendorf *et al*. 2001). Hybridization has played a major role in the origins of diversity among species that we see today. For example, hundreds of new cichlid fishes resulted from natural hybridization between two divergent lineages of cichlids (Meier *et al*. 2017). Although these fish provide a clear example of hybridization resulting in speciation, often the effects of hybridization on biodiversity are unclear or equivocal. We struggle with defining species when boundaries are blurred and hybridization is an ongoing event. These interbreeding populations can be subjected to demographic and genetic swamping where individuals are being either physically or genetically replaced by hybrids (Rhymer & Simberloff 1996, Wolf *et al*. 2001, Todesco *et al*. 2016). Populations that exhibit ranges of ancestry in individuals make defining species difficult. Often, we are unable to define hybridized individuals as a specific species because they have ancestry coming from two or more different species. This inability to concisely define an individual creates space for authors to use their own definition of hybrids, leading to confusion and lack of meaning behind given definitions.

The term hybrid swarm has been regularly used in studies of hybridization. It was originally defined in 1926 as a dynamic group with hybrids that have intermediate traits which may be biased toward one parental species (Cockayne & Allan 1926). However, when we look at the way this term has been used over time, we see that its use is quite ambiguous. “Hybrid swarm” appears to be used in various different ways and with multiple different meanings. Our goal in this study was to better understand what most authors mean by hybrid swarm, quantify how consistent usage is across taxonomic groups, and provide a perspective on the advisability of continued use of this term. In this review we summarized how the term hybrid swarm is used in the literature by investigating species studied, methods used for identification of hybrids (genetic versus phenotypic), generations of hybridization described, and number of species involved. We conducted a comprehensive review of published literature using the term hybrid swarm, analyzed our compiled data, and drew data-driven conclusions which highlight the equivocal use of the term hybrid swarm.

### 3 Methods

### 3.1 Data collection

We collected scientific articles for this review using a standardized search method in R. Using four packages, `easyPubMed` (Fantini 2019, version 2.13), `remotes` (Csárdi *et al*. 2021, version 2.4.0), `litsearchr` (Grames *et al*. 2019, version 1.0.0), and `plyr` (Wickham 2011, version 1.8.6), we created a custom script that searched and pulled publications from the PubMed database. Search terms “hybrid swarm” and “biology” were used to provide bounds for the PubMed database search. After the initial search, our results contained many articles that were not pertinent to the scope of this review. These articles contained the word “particle” in addition to “hybrid swarm” and “biology”. Therefore, we also we modified our query terms to not include the word “particle”. We conducted the search to pull 74 articles at a time, resulting in a representative sample of papers relevant to our objectives. Key information such as paper title, publishing journal, authors, date published, and paper abstract was imported and written to a .csv file. This information was then used to find and download these papers in full. Two separate searches were done approximately two years apart for this project. The first search was on February 8th, 2021 and the second on March 27th, 2023.

All scientific articles pulled were read and assessed for fit in the scope of this review; we classified each paper as either relevant or not to hybridization in an evolutionary biology context and either describing an ongoing hybrid swarm or not; in many cases, a “no” in the “ongoing hybrid swarm” category meant the authors stated that the system was either descended from or could become a hybrid swarm, rather than actively existing as one during the time of study. If we determined that the paper was relevant, two different people read each paper and data were manually collected on the following: type of evidence for hybridization (genetic, phenotypic, etc.), taxonomic group (e.g., bird, fish, plant), genus, number of species involved in hybridization, number of individuals sampled, number of genetic loci employed (or NA if no genetic data were used), number of sampling locations, years of hybridization, genetic marker type (or NA if no genetic data were used), number of generations of hybrids (F1, F2, or beyond), number of generations of backcrossing, and finally, binary responses on whether anthropogenic disturbances, non-anthropogenic disturbances, or species introductions were present in the system and whether data was available for the study. We then cleaned the data by merging the data from both readers into one entry per paper and manually evaluating and reconciling differences between the two readers’ data entries.

### 3.2 Data analysis

We created a timeline of some of the major events relating to the use of hybrid swarm and other major events related to genetics (Fig. 1). We also plotted the frequency of use of hybrid swarm for all relevant papers, coloured by those which describe active hybrid swarms (Fig. 2) and summary statistics on the cleaned data (Figs. 3, 4). Finally, we created a Sankey plot depicting outcomes of hybridization for all relevant papers Fig. 5). We used R Statistical Software (Team” 2021, v4.1.2) to create all figures, and used the packages `tidyverse` (Wickham *et al*. 2019, version 1.3.1), `RColorBrewer` (Neuwirth 2014, version 1.1.2), `scales` (Wickham *et al*. 2022, version 1.2.1), `patchwork` (Pedersen 2023, version 1.1.3), and `ggsankey` (Sjoberg 2021).

**Figure 1.**
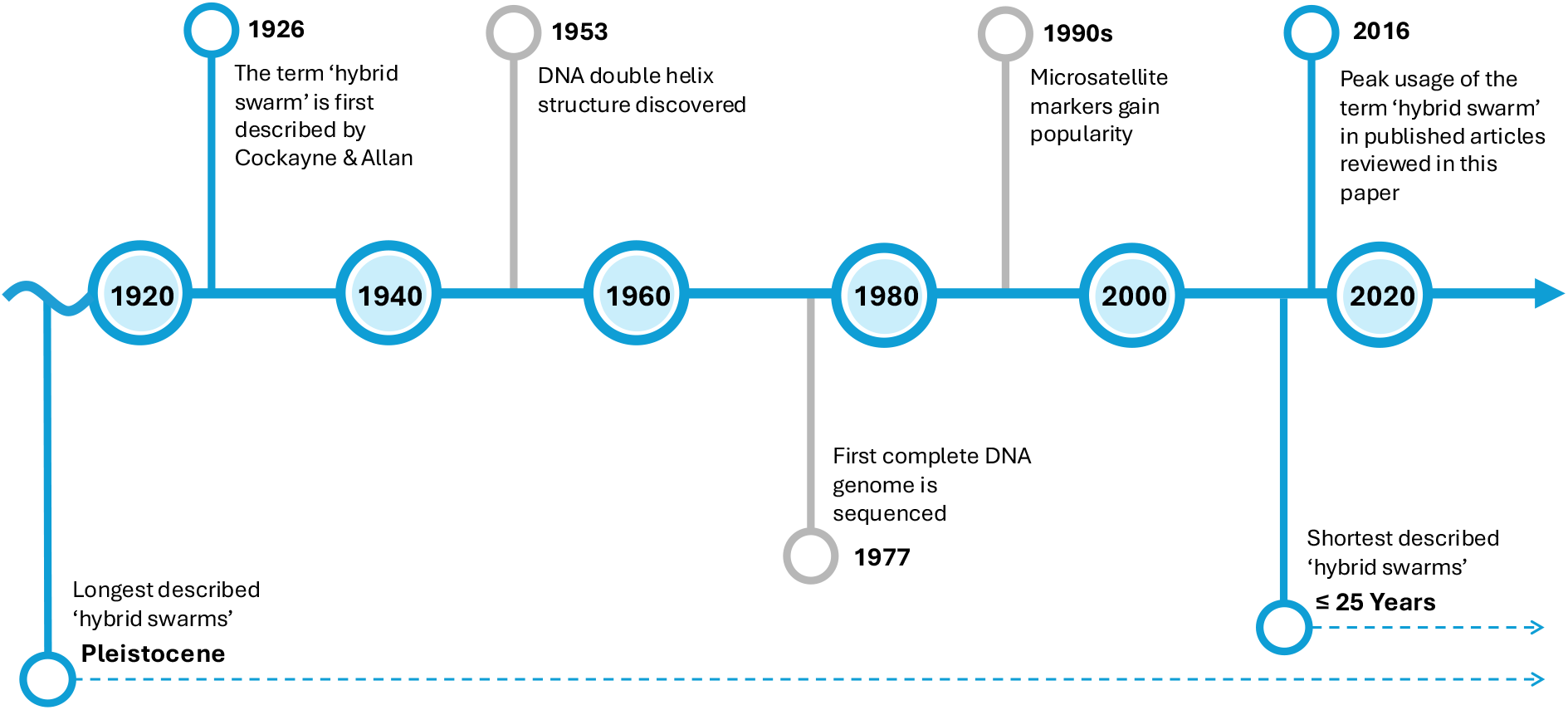
Timeline depicting usage of the term “hybrid swarm” (blue) and important historic events (grey) surrounding genomic research. Figure adapted from Martínez-Pèrez et al. (2022).

## 4 Results

From a total of 148 papers, 111 were unique between the two pulls. From these, we classified 67 as relevant to hybridization in an evolutionary biology sense and 49 as including active hybrid swarms. The first usage of hybrid swarm within the papers pulled by our search was in 1989 (Fig. 2). The search algorithm prioritized more recent studies, so this is not an exhaustive list of papers using the term “hybrid swarm”, but this focus on the past 35 years of publications suits our goal of assessing current usage of hybrid swarm in the field. There is a peak in usage of hybrid swarm around 2016-2019 (Fig. 1, 2).

**Figure 2.**
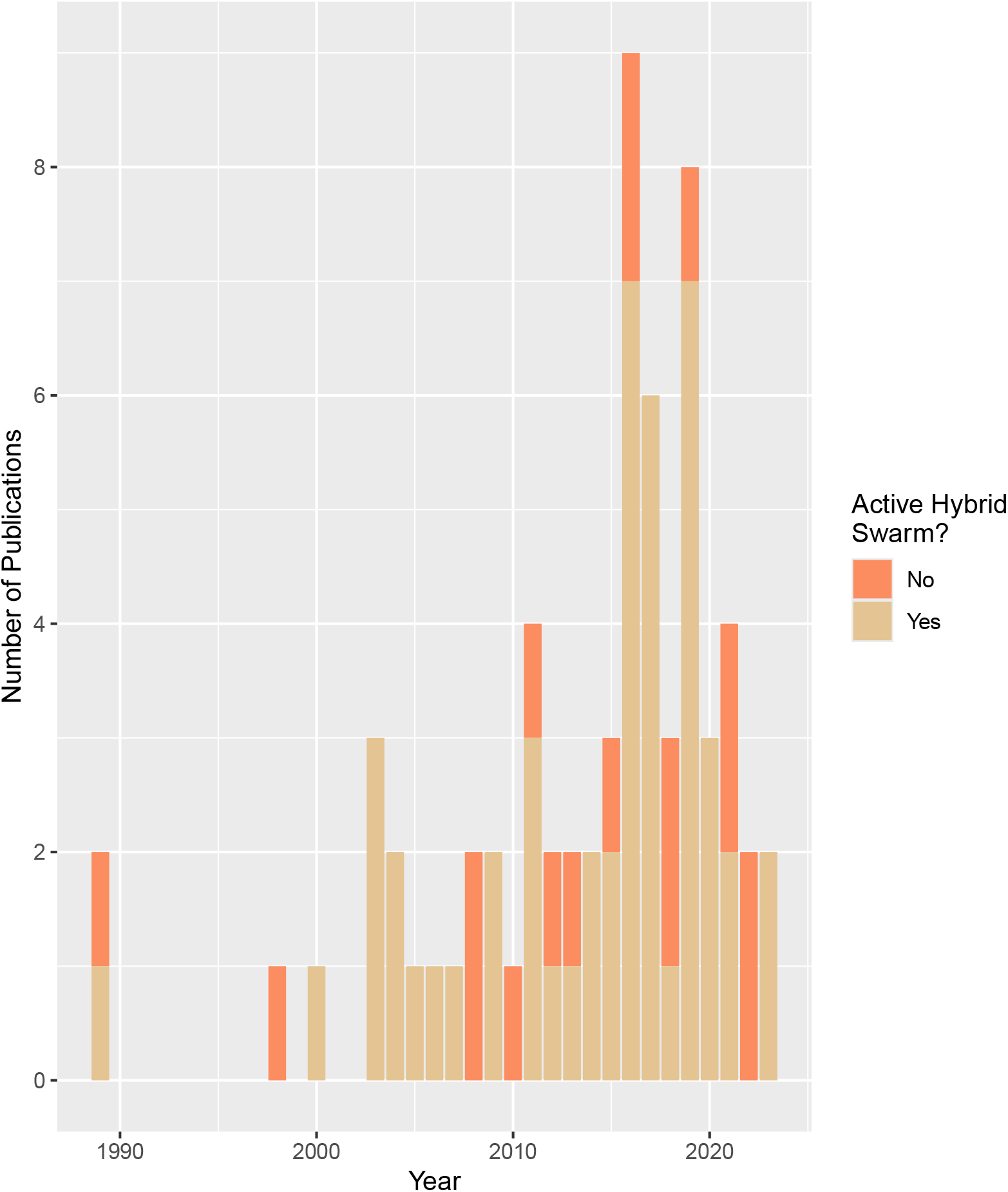
Frequency of the usage of the term “hybrid swarm” over time for all 67 relevant papers.

Plants were by far the most commonly studied group to be referred to as a hybrid swarm, while amphibians, fungi, and protozoa were the least common (Fig. 3A). Two-species hybrid systems were the most commonly studied systems (Fig. 3B). Single-species systems were the second most popular, due to many papers on hybridization between sub-species. There was quite a bit of variation in the number of individuals per study, though most were 500 or fewer (Fig. 3C,D). A similar trend was seen with the number of sampling sites and loci used in a study, where there were some outliers with many, but most studies used very few (fewer than 25 or 100, respectively) (Fig. 3E-H).

**Figure 3.**
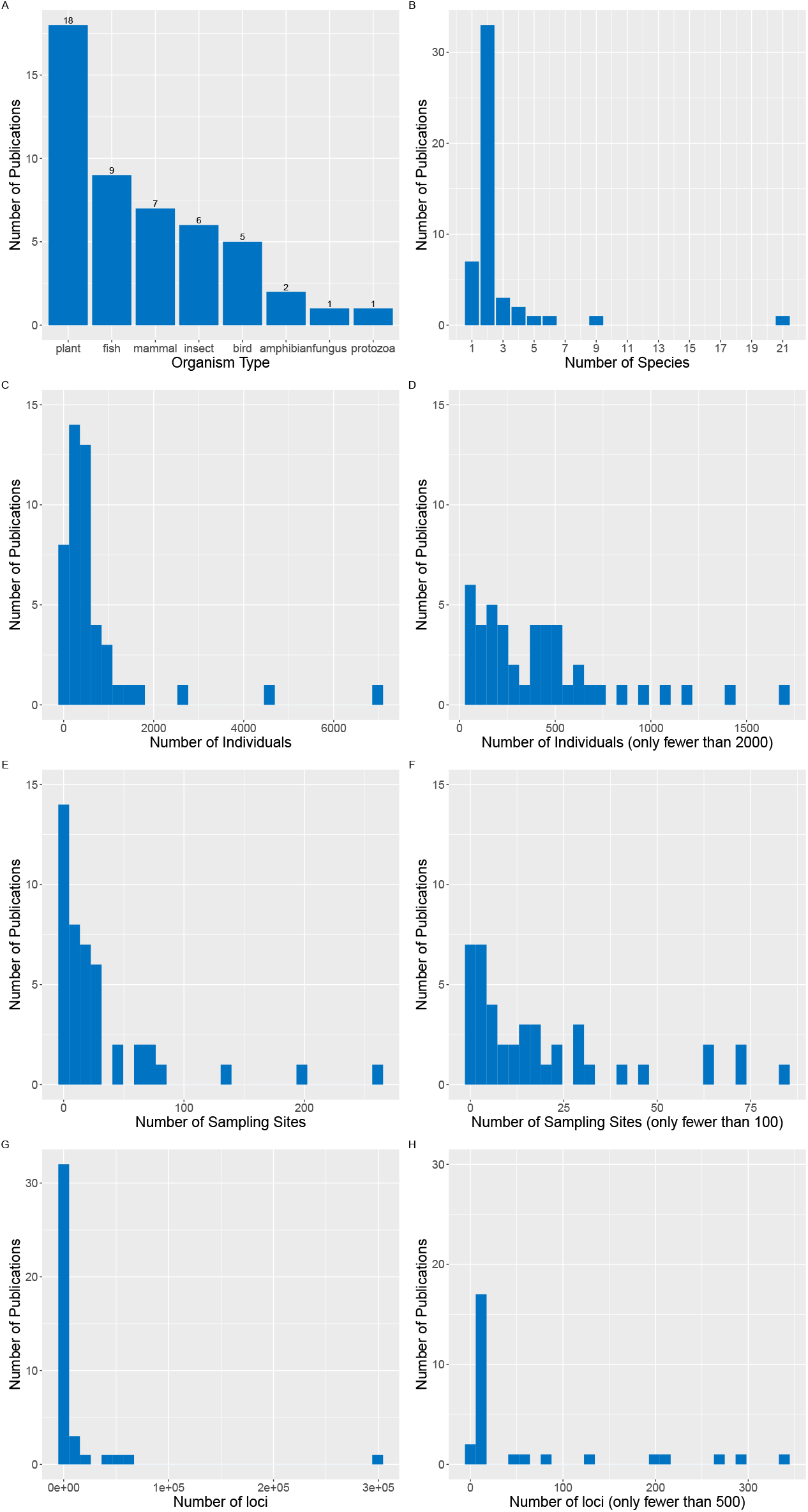
Summary statistics on the systems studied in all 49 papers which described hybrid swarms. A. Number of publications per type of organism. B. Number of publications per number of species studied. C. Number of publications per number of individuals used. Number of publications per number of individuals used, for publications with fewer than 2000 individuals. E. Number of publications per number of sampling sites used. F. Number of publications per number of sampling sites used, for publications with fewer than 100 sites. G. Number of publications per number of loci used. H. Number of publications per number of loci, for publications with fewer than 500 loci.

SNPs were more frequently used to describe hybrid swarms in recent years, while microsatellite marker usage is becoming less frequent (Fig. 4A,B). Genetic evidence and phenotypic evidence usage have both been fairly stable over the years, with genetic being used more overall (Fig. 4C,D). Data from publications describing hybrid swarms were published more frequently in recent years (Fig. 4E). Disturbance is much more common among systems with fish that are described as hybrid swarms than amongst other systems, while the opposite is true for plants (Fig. 4F).

**Figure 4.**
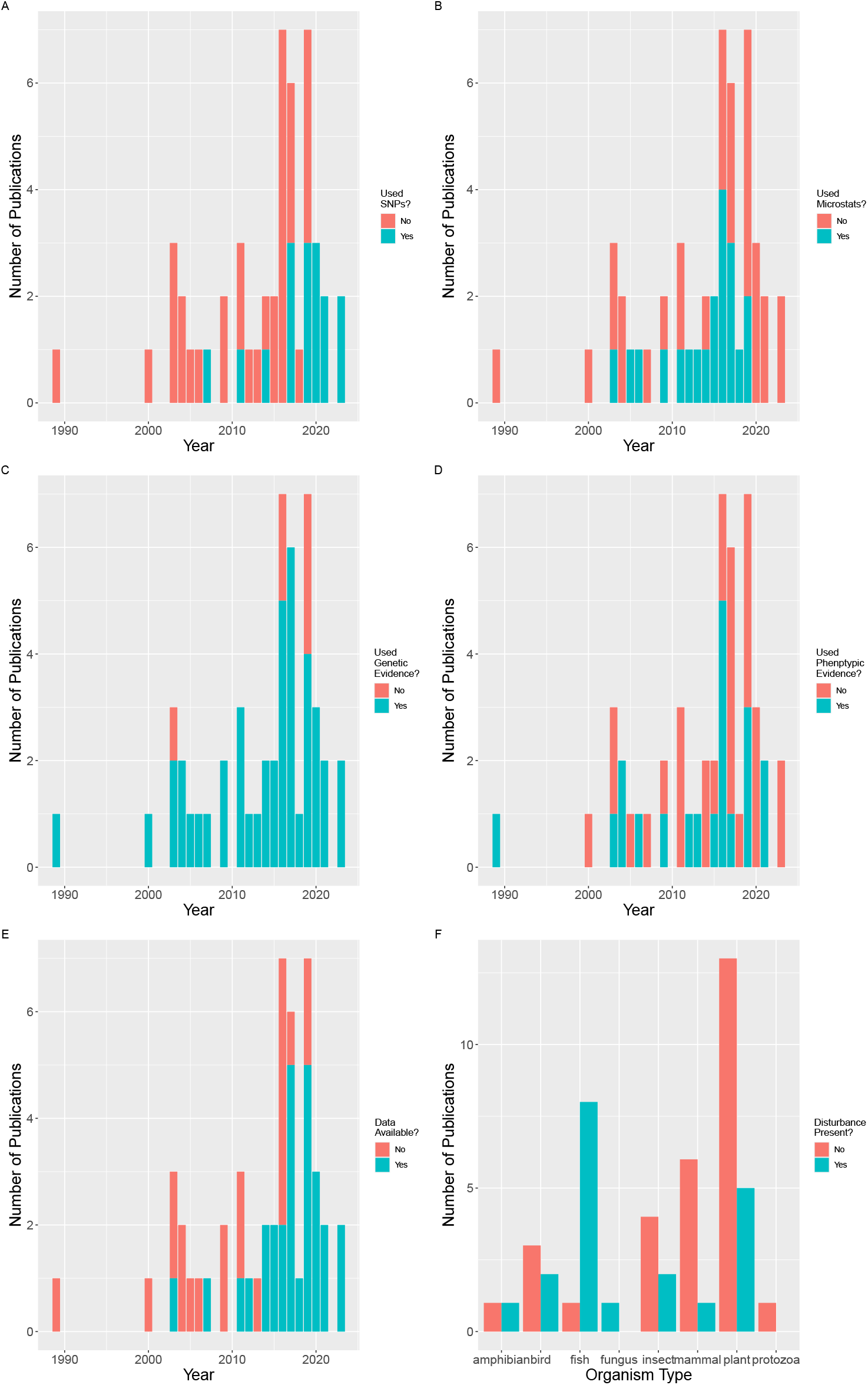
Summary statistics on types of data used in all 49 papers which described hybrid swarms. A. The number of publications which used SNPs, by year. B. The number of publications which used micro-satellite markers, by year. C. The number of publications which used genetic evidence, by year. D. The number of publications which used phenotypic data, by year. (Subfigures C and D are not mutually exclusive – some publications used both or neither) E. The number of publications which have published data available, by year. F. The number of publications which had some form of disturbance present in their study system, by organism type.

Most of the active hybrid swarm systems (31/49) had F1s present (Fig. 5). Two common outcomes were either having all 3 hybrid types present (F1s, F2 or later generations, and backcrosses) or having none (18/49 and 17/49, respectively). About half of all papers (25/49) recorded having backcrosses present. A small number (4/49) papers had F1s present but no other hybrids, and only one system had F2s or later generation hybrids without also having F1s.

**Figure 5.**
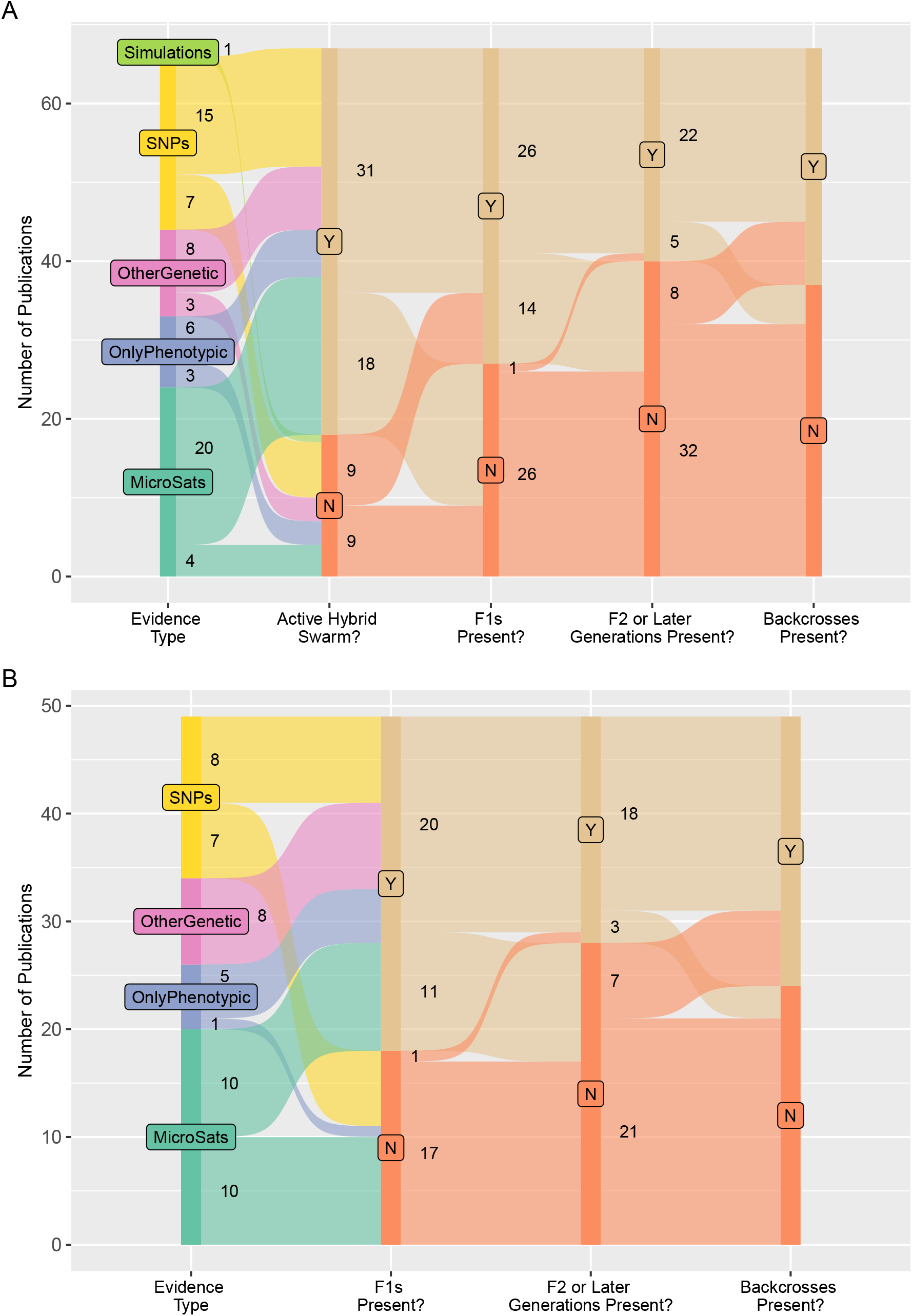
Sankey plot depicting hybrid outcomes. Evidence type describes the type of evidence that a paper used to detect hybrids in a system; papers that used a type of genetic marker may have also used non-genetic evidence as well. A. All 67 relevant papers. B. Only the 49 papers depicting active hybrid swarms.

## 5. Discussion

One of the first usages of the term hybrid swarm was by Cockayne & Allan (1926). They stated that a hybrid swarm is dynamic, unlike a species, which is static, and that there are hybrids that look more or less intermediate, often biased towards one parental species. However, what exactly constitutes a hybrid swarm remains elusive even after our literature review and data analysis. The papers we reviewed described many very different and distinct evolutionary scenarios as hybrid swarms. Many of the papers we reviewed stated that the presence of a large number of backcrosses or exclusively backcrosses in a hybrid zone are characteristics of a hybrid swarm (Anttila *et al*. 2000, Glotzbecker *et al*. 2016, Haines *et al*. 2019, Jacquemyn *et al*. 2016, Keim *et al*. 1989, Lavretsky *et al*. 2019, Opiro *et al*. 2017, Stemshorn *et al*. 2011, Tihon *et al*. 2017), but we found much evidence to the contrary in our analysis as only about half of papers with active hybrid swarms described the presence of backcrosses (Fig. 5). Four of the papers we reviewed showed F1s present but no later generation hybrids or backcrosses (Campoy *et al*. 2019, Mallet *et al*. 1998, Marczewski *et al*. 2016, Van Der Sluijs *et al*. 2008), while other papers stated that indeed continual hybrid generations and lack of backcrossing with parental species is actually characteristic of a hybrid swarm (Li *et al*. 2016, Lamont *et al*. 2003, Latch *et al*. 2011). In addition, more papers said both backcrossing with parentals and mating between hybrids are found in hybrid swarms (Glotzbecker *et al*. 2016, for example). Seventeen papers were actually classified as having no hybrids or backcrosses present at all, indicating that they were not able to discern between hybrid types (Anttila *et al*. 2000, Taylor *et al*. 2006, Mende *et al*. 2016, as examples). Studies on hybridization that used SNP data had more power to accurately describe hybridization outcomes (McFarlane & Pemberton 2019), potentially eliminating the need for a vague, catch-all term like hybrid swarm. Many of the papers that we reviewed used the term hybrid swarm when they were unable to confidently differentiate hybrid generations or backcrosses from one another. Thus, while an initial impression from reading these papers may be that a hybrid swarm can be quite clearly defined as the presence of many backcrosses in a system, we have found that not to be the case. From our data, we found that most of these hybrid swarms had F1s present, some had continued hybrid generations, and about half had backcrosses present (Fig. 5); overall, there was no clear consensus as to what hybrid outcomes are present in a hybrid swarm.

In our review of the term in recent decades, we saw a slow increase in usage from the 1990s until hitting a peak around 2016 to 2019, after which usage declined (Fig. 2). We attribute this increase in usage to the increase in studies likely being published on hybridization in wild systems, as the reduction in cost of sequencing a large number of individuals has decreased over the last two decades and the usage of SNP data has become more popular (Davey *et al*. 2011, Lou *et al*. 2021, Narum *et al*. 2013). The usage of SNPs increased and the usage of microsatellite markers decreased as well (Fig. 4A, B).

The term hybrid swarm has been used to describe hybridization dynamics in systems with a great diversity in terms of the types of taxa and the number of loci, individuals, and sampling sites used (Fig. 3ACDEFGH). However, the vast majority of the papers that we examined were studies done on 2-species systems (Fig. 3B). This exemplifies how multi species dynamics are not as well studied and again supports the idea that the term hybrid swarm is complicit with an oversimplification of hybridization in wild systems.

There was also no consensus on how stability or duration may define hybrid swarm. Fitzpatrick & Schaffer (2007) described a system of hybridization between two species of amphibians subject to anthropogenic disturbance; they used both the term “hybrid zone” and “hybrid swarm”, and implied that hybrid zones are areas of stable hybridization, while hybrid swarms are expanding and threatening to cause genetic and/or demographic swamping of the species. We have shown that hybrid swarm was used to describe systems with or without disturbance (Fig. 4F). Additionally, the term was used to describe hybrid zones that have existed for vastly different time periods, including some present for 25 years or less at the time of publication (Anttila *et al*. 2000, Erickson *et al*. 2020, Glotzbecker *et al*. 2016, Kirk *et al*. 2005, Knutson *et al*. 2019, Rangel *et al*. 2016, Ward *et al*. 2012) as well as hybrid zones which have potentially been in existence since the end of the Pleistocene (Li *et al*. 2016, McDevitt *et al*. 2009) or earlier (Dupuis & Sperling 2015) (Fig. 1). It is clear that the term has been used without regard to the stability of the hybrid zone or duration of its existence.

Based on our synthesis of published papers, we argue that these so-called hybrid swarms are simply instances of hybridization and that there is no need for the specific term hybrid swarm. Instead, we advocate for more precisely describing dynamics of hybridization in a system, as we can now do with genomic data. Crystal et al. (2016) stated that hybrid swarms are “characterized by high phenotypic variability among individuals with extremely diverse genotypes.” This is corroborated by Dejaco et al. (2016) who also stated that “when two distantly related genomes fuse, hybrid swarms consisting of lineages with different admixture proportions can emerge… hybrid swarms can act as cradle of novel, highly adapted genotypes”. While this is true, it is not a novel phenomenon that warrants a name independent of “hybridization”. Many studies have shown that hybridization can cause novel phenotypes, unique from that of either parental species (e.g. Hegarty *et al*. 2008, Rieseberg *et al*. 1999).

Using the data generated from our literature search, we showed that there is little consensus on what hybrid outcomes constitute a hybrid swarm (Fig. 5). Moreover, we were unable to connect the use of hybrid swarm with specific hybridization scenarios. We had originally hypothesized that hybridization as a result of disturbance might be more frequently referred to as a hybrid swarm, but there is no evidence that the term is more commonly used in disturbed areas (Fig. 4F). There is also no evidence that the use of the term hybrid swarm is related to how long hybridization has been occurring (Fig. 1).

Fitness is also not a key component of hybridization beign described as a hybrid swarm. In LeRoy et al. (2016), the authors describe a hybrid swarm as “a population of hybrids at least as vigorous as parental genotypes”. With this description, the authors are referring to the hybrid populations of their studied pathogen being as or more aggressive than the parental populations. However, the majority of papers on hybridization do not have a direct measure of fitness by which they can quantify hybrids as part of a hybrid swarm. The authors go on to say that hybrid swarms are characterized by high genotypic and phenotypic variances because they are produced by admixture of divergent populations.” (Leroy *et al*. 2016). However, this description is not unique to these so-called “hybrid swarms” – this same description could be used for any population in which hybrids exist. Hybridization is the mating between divergent populations (Stebbins 1959).

Often, we have found that the context surrounding usage of the term hybrid swarm is concerned with the evolutionary outcomes of the hybridization. In these cases, it seems that the terms demographic swamping and genetic swamping would be better employed. Todesco et al. (2016) explained these terms quite eloquently; in essence, demographic swamping is when the waste of reproductive efforts between two species on the production of sterile F1 hybrids causes one species to go extinct, while genetic swamping is continual mating between two species and their fertile offspring, causing the replacement of all “pure” parental individuals with hybrids or backcrosses. We noted at least 3 cases in the papers we reviewed, that would benefit directly from the usage of this clearer terminology (Liou & Price 1994, Lavretsky *et al*. 2019, Wells *et al*. 2019). In contrast, Pinto et al. (2005) somewhat bucked this trend, stating that “the degree of mixing between hybridizing forms may range from formation of a hybrid swarm to genetic assimilation of one form by the other.” This definition seemingly distinguishes the formation of a hybrid swarm from genetic swamping. However, if a hybrid zone is able to be maintained without an overall loss of biodiversity via demographic or genetic swamping, this would be considered a stable hybrid zone (Todesco *et al*. 2016). All in all, we continue to find that the usage of the term hybrid swarm is not the ideal terminology to use in any context.

Contrarily, one aspect of a hybrid swarm was quite consistent across definitions – how hybrid swarms are formed. Many of the papers we reviewed described hybrid swarm formation as a lack of strong selection against or assortative mating between parental populations and/or hybrids (Haines *et al*. 2019, Hasselman *et al*. 2014, Mallet *et al*. 1998, Natola *et al*. 2022, Nielsen *et al*. 2003, Van Der Sluijs *et al*. 2008). But again, this is not a feature unique to a phenomenon that can be succinctly described with the term “hybrid swarm”; hybridization often occurs when there is a lack of strong selection against or assortative mating between populations upon secondary contact (Todesco *et al*. 2016).

We argue that the terms “hybrid zone” and “hybridization” can be used in replace of “hybrid swarm” in most cases. We observed these two terms used interchangeably in many studies already (for example Fitzpatrick & Shaffer 2007, McDevitt *et al*. 2009), and have also found a description stating that hybrid swarms develop within hybrid zones, again indicating that a hybrid swarm is somehow a unique form of hybridization (Glotzbecker *et al*. 2016). Here, we define that “hybrid zone” signifies an area in which hybridization is occurring and hybrid offspring are being produced. Hybridization outcomes within the hybrid zone may then be described with terms including (but not limited to) “stable hybrid zone”, “demographic swamping”, and/or “genetic swamping” (Todesco *et al*. 2016), depending the types of hybrid observed and quantified using genomic and phenotypic data. Overall, we have found the term “hybrid swarm” to be ambiguous and used in many different contexts. We have found it to not represent any unique concept that cannot be signified by another term or better described using more precise language.

This review was prompted by the large variety of terminology used to describe hybridization. As new terminology is developed in this ever growing field, it is necessary to reflect upon and optimize what is being used, to be able to effectively describe research and succinctly convey findings. We hope that this body of work will provoke thoughtful discussion and consideration of the terminology best suited to describe hybridization.

## Acknowledgements

J.N. Campbell was supported by Species Conservation Trust Fund Project Number SCA20BXYZ from Colorado Parks and Wildlife. N.C. Lewis was supported from a USRA award from NSERC at the University of Guelph. A.V. Meuser was supported in part by the Canada First Research Excellence Program, through the Food From Thought program at the University of Guelph. We would like to thank Kayleigh Paisley-Rush, Matthew Mervyn, and Teaghan Frauley, who contributed to early stages of the project as undergraduate researchers. This manuscript was improved by comments from Katherine Drotos and members of the Mandeville Lab at Northern Michigan University.

## 6 Data Availability

`R` scripts used for the PubMed search and creating figures can be found at https://github.com/amanda-meuser/Hybrid_Swarm_Paper/. Raw data files will be published on the Dryad data repository upon the acceptance of this manuscript.

## Notes

### Competing Interest Statement

The authors have declared no competing interest.

